# DNA capture and loop extrusion dynamics by cohesin-NIPBL

**DOI:** 10.1101/2022.08.18.504320

**Authors:** Parminder Kaur, Zhubing Shi, Xiaotong Lu, Hongshan Zhang, Ilya J. Finkelstein, Yizhi Jane Tao, Hongtao Yu, Hong Wang

## Abstract

3D chromatin organization plays a critical role in regulating gene expression, DNA replication, recombination, and repair. While initially discovered for its role in sister chromatid cohesion, emerging evidence suggests that the cohesin complex (SMC1, SMC3, RAD21, and SA1/SA2), facilitated by NIPBL, mediates topologically associating domains (TADs) and chromatin loops through DNA loop extrusion. However, information on how conformational changes of cohesin-NIPBL drive its loading onto DNA, initiation, and growth of DNA loops is still lacking. Using high-speed AFM (HS-AFM) imaging, we show that cohesin-NIPBL captures DNA through arm extension, followed by transfer of DNA to its globular domain and DNA loop initiation independent of ATPase hydrolysis. Additional shorter protrusions (feet) from cohesin-NIPBL transiently bind to DNA, facilitating its loading onto DNA. Furthermore, HS-AFM imaging reveals distinct forward and reverse DNA loop extrusion steps by cohesin-NIPBL. These results provide critical missing links in our understanding of DNA binding and loop extrusion by cohesin-NIPBL.

## INTRODUCTION

Large-scale spatial segregation of open and closed chromatin compartments and topologically associating domains (TADs), sub-TADs, and loops fold the genome in interphase (1-5). TADs that contain continuous regions of enriched contact frequencies play essential roles in the timing of DNA replication (6), regulation of enhancer-promoter contacts, gene expression, DNA repair, and V(D)J recombination (7-10). The structural maintenance of chromosomes (SMC) protein family, including cohesin and condensin complexes, play critical roles in 3D chromatin organization in all living organisms (11-13). The core cohesin complex includes SMC1, SMC3, RAD21^Scc1^, and SA1/SA2 ^Scc3^ (human^yeast^, **Figure 1a**). SMC proteins (SMC1 and SMC3) form long antiparallel coiled coils (arms), each with a dimerization (hinge) domain at one end and an ATP-binding cassette (ABC)-type ATPase (head) domain at the other. RAD21^Scc1^ interconnects the head domains. In addition, SA1 and SA2 directly interact with the CCCTC-binding factor (CTCF), a ubiquitous zinc-finger (ZF) protein that specifically localizes to CTCF binding sites (CBS) along the genome (14). Though initially identified as an essential complex to hold sister chromatids together (15), numerous studies demonstrated that cohesin is also crucial in mediating 3D chromatin organization during interphase (16-21). Greater than 80% of long-range looping interactions are mediated by some combinations of cohesin, CTCF, and the Mediator complex. Cohesin and CTCF are enriched at TAD boundaries and corner peaks that indicate strong interactions at TAD borders (2,5). Furthermore, NIPBL significantly stimulates DNA binding and ATPase activities of cohesin (22). RAD21 or NIPBL depletion leads to significantly reduced TADs and corner peaks.

**Figure 1.**
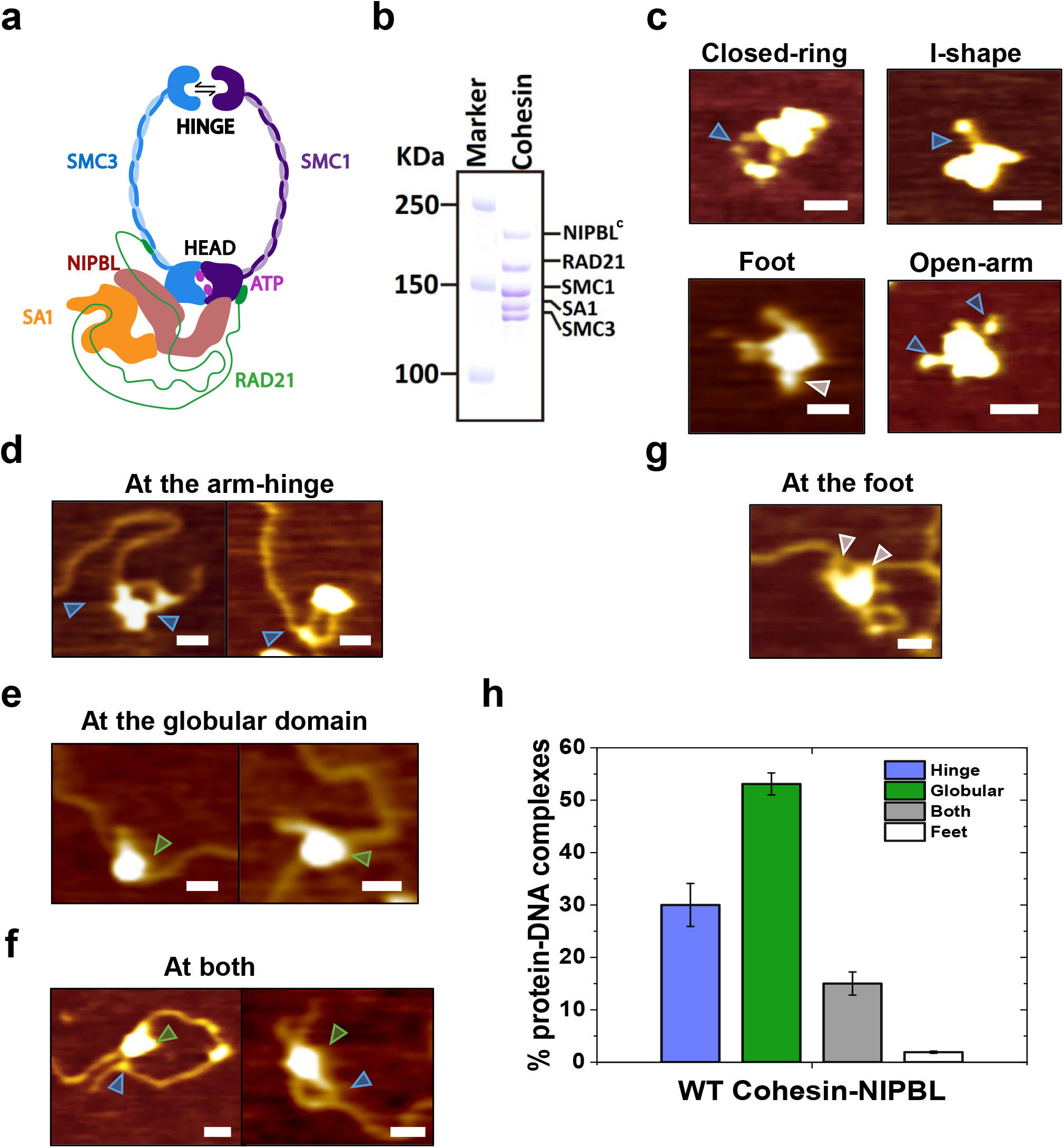
AFM imaging in air shows diverse conformations of WT cohesin-NIPBL alone and DNA binding by cohesin-NIPBL. (**a**) Schematic representation of cohesin-NIPBL based on the cryo-EM structure. (**b**) SDS-PAGE of cohesin^SA1^-NIPBL^c^ showing individual subunits. (**c**) Example AFM images showing cohesin^SA1^-NIPBL^c^ alone. (**d** to **g**) Example AFM images of cohesin^SA1^-NIPBL^c^ binding DNA (5.19 kb, + 2.5 mM ATP) at the arm-hinge (**d**), globular domain (**e**), both the arm-hinge and globular domains (**f**), and the foot (**g**). XY scale bar = 50 nm. Blue arrow: arm; green arrow: globular domain; gray arrow: foot. (**g**) Percentages of cohesin^SA1^-NIPBL^c^-DNA complexes with the arm-hinge, globular, both arm-hinge and globular domains, and foot binding to DNA. N=105 protein-DNA complexes. Error bars: SD. Two independent experiments.

A large body of literature supports a model that cohesin-NIPBL mediates TAD and chromatin loop formation through DNA loop extrusion (23-26). The DNA loop extrusion model posits that cohesin creates DNA loops by actively extruding DNA until they are stabilized by CTCF bound at converging CBS (27,28). Importantly, single-molecule fluorescence imaging demonstrated that cohesin-NIPBL is capable of DNA loop extrusion in an ATPase-dependent manner (22,23). One emerging consensus is that cohesin-NIPBL contains multiple DNA binding sites (25). Previous high-speed AFM (HS-AFM) imaging also showed that cohesin and condensin alone are highly dynamic and capable of significant conformational changes, including SMC ring opening and closing, alignment of the SMC arms, elbow bending, and SMC head engagement and disengagement (25,29-33). However, HS-AFM has not been applied to study cohesin with DNA (25). Meanwhile, observations from single-molecule fluorescence imaging did not provide information on protein conformational changes that drive DNA binding and loop extrusion and could miss intermediate DNA loop extrusion steps by cohesin (22).

Hence, because of technical challenges in studying dynamic multi-subunit cohesin-NIPBL complexes, the mechanism of DNA binding and loop extrusion by cohesin is still under intense debate (25,26,33-36). Several key questions remain unanswered regarding DNA binding and loop extrusion by cohesin-NIPBL, such as: How does cohesin-NIPBL load onto DNA and initiate a DNA loop? What are the DNA loop extrusion step sizes and associated cohesin-NIPBL conformational changes that drive DNA binding and loop extrusion?

Here, we applied traditional AFM imaging in air and HS-AFM imaging in liquids to reveal the structure and dynamics of cohesin-NIPBL mediated DNA binding and loop extrusion. These studies show that cohesin-NIPBL uses arm extension to capture DNA and initiate DNA loops independent of ATPase hydrolysis. Surprisingly, foot-like protrusions on cohesin-NIPBL can transiently bind to DNA and facilitate DNA segment capture by the arm-hinge domain. Furthermore, HS-AFM imaging reveals distinct forward and reverse DNA loop extrusion steps. These results shed new light on the cohesin-mediated DNA loop extrusion mechanism and provide new directions for future investigation of diverse biological functions of cohesin.

## MATERIALS AND METHODS

### Protein purification

WT and SMC1A-E1157Q/SMC3-E1144Q (EQ) ATPase mutant cohesin^SA1^-NIPBL^c^ (SA1 containing cohesin with the C-terminal HEAT repeat domain of NIPBL) were purified according to protocols published previously, which involved mixing purified subcomplex containing SMC1, SMC3, RAD21, and NIPBL^c^ with separately purified SA1 (22). Purification of N-terminal His_6_-tagged full-length SA2 and SA2 fragments (1-301, 1-450, 451-1051, and 1052-1231 AAs) was described previously (37-40).

### DNA substrates

pG5E4-5S plasmid (5190 bp, a gift from the Williams lab at UNC-Chapel Hill) was linearized using NdeI restriction enzyme (NEB) and purified using the Qiagen PCR purification kit. The 45 bp duplex DNA was prepared as described previously (39).

### AFM Imaging in air

Purified linear dsDNA (6 nM, 5190 bp) was incubated with WT or ATPase mutant cohesin^SA1^-NIPBL^c^ (30 nM) in Cohesin Buffer (40 mM Tris pH 8, 50 mM NaCl, 2 mM MgCl_2_, 1 mM DTT) either without or with ATP (2.5 mM) for 1 min at room temperature. All samples were diluted 16-fold in AFM Imaging Buffer (20 mM HEPES pH 7.6, 100 mM NaCl, and 10 mM Mg (C_2_H_3_O_2_)_2_) and immediately deposited onto a freshly cleaved mica surface. The deposited samples were washed with deionized water and dried under nitrogen gas streams before AFM imaging. AFM imaging in air was carried out using the AC mode on an MFP-3D-Bio AFM (Asylum Research, Oxford Instruments) with Pointprobe PPP-FMR probes (Nanosensors, spring constants at ∼2.8 N/m). All images were captured at scan sizes of 1 × 1 μm^2^ to 3 × 3 μm^2^, a scan rate of 1–2 Hz, and a resolution of 512 × 512 pixels. DNA bending angle analysis was done using Image J software.

### High-speed atomic force microscopy (HS-AFM) imaging in liquids

WT or ATPase mutant cohesin^SA1^-NIPBL^c^ (30 nM) was incubated with the linear dsDNA substrate (3 nM) in Cohesin Buffer for 1 min at room temperature, followed by a 1 min incubation with ATP (4 mM). The incubated sample was diluted 20-fold in Cohesin Buffer and deposited onto a freshly prepared 1-(3-Aminopropyl)silatrane (APS)-treated mica surface (41). APS was synthesized in-house to ensure high purity. The protein-DNA sample was further incubated on the APS-mica surface for 2 min, followed by washing with Cohesin Buffer (500 μl). The washed sample was scanned in Cohesin Buffer containing ATP on either a Cypher VRS AFM (Asylum Research) using BioLever fast (AC10DS) cantilevers or JPK NanoWizard 4 using USC-F0.3-k0.3 cantilevers. For Cypher VRS, we used BlueDrive Photothermal Excitation to drive the cantilever. The images were scanned at 0.4-2.3 frames/s and analyzed using either Asylum or JPK image analysis software.

### Fluorescence anisotropy

His_6_-tagged full-length SA2 and SA2 fragments in DNA Binding Buffer (20 mM HEPES, pH 7.5, 0.1 mM MgCl_2_, 0.5 mM DTT, 100 mM KCl) were titrated into the binding solution containing fluorescein-labeled DNA substrates (6 nM, 45 bp) using a Tecan Spark Multimode plate reader (Tecan Group Ltd) (40). The data obtained from fluorescence anisotropy were analyzed by using the equation *P* = ((*P*_bound_− *P*_free_)[protein]*/*(*K*d + [protein])) + *P*_free_, where *P* is the polarization measured at a given total protein concentration, *P*_free_ is the initial polarization of fluorescein-labeled DNA without protein binding, *P*_bound_ is the maximum polarization of DNA due to binding of proteins, and [protein] is the total protein concentration. The average equilibrium dissociation constant (K_d_) was based on two to three measurements.

### Statistical Analysis

Data from AFM imaging in air and liquids were pooled from at least two to three independent experiments. Statistical analysis was carryout out using OriginPro (OriginLab). Unless stated otherwise, the error bars represent SD. The P-value was calculated by Student’s t-test, and the statistically significant level was set at p<0.05. *P < 0.05, **P < 0.01, ***P < 0.001, ****P < 0.0001.

## RESULTS

### DNA binding and initiation of DNA loops by cohesin-NIPBL

Recent studies demonstrated that cohesin-NIPBL contains multiple DNA binding sites on the interface between SMC1 and SMC3 hinges, SMC heads, SA1/SA2 (38), and NIPBL (25). These DNA binding sites are essential for DNA loop extrusion (25). Despite these new discoveries, our understanding of how each DNA binding domain on cohesin-NIPBL contributes to cohesin loading onto DNA is limited. To directly address this question, we purified WT (**Figure 1b**) and ATPase-deficient SMC1A-E1157Q/SMC3-E1144Q (EQ) cohesin^SA1^-NIPBL^c^ (SA1 containing cohesin with the C-terminal HEAT repeat domain of NIPBL) mutant (22,42). We applied AFM imaging in air and HS-AFM imaging in liquids (43) to investigate the structure and dynamics of cohesin^SA1^-NIPBL^c^ alone and in complex with DNA. Consistent with the previous literature (25), AFM images of the cohesin^SA1^-NIPBL^c^ collected in air (+ 2.5 mM ATP, **Figure 1c**) showed monomers with their SMC arms (blue arrows, **Figure 1c**) distinguishable from the globular domain (including SMC heads, RAD21, SA1, and NIPBL^c^). Based on their distinct arm features, cohesin^SA1^-NIPBL^c^ monomers (N_total_=127) can be categorized into several classes (**Figure 1c**), including closed-ring (18.1%), I-shape with aligned SMC arms (23.8%), two open arms (21.3%), and those unclassifiable (36.8%). Unexpectedly, in addition to arms, some cohesin^SA1^-NIPBL^c^ molecules (20.5%) showed either one or two small protrusions (feet) at the bottom of the globular domain (**Figure 1c**).

To study DNA binding by cohesin-NIPBL, we first employed AFM imaging in air to examine samples of cohesin^SA1^-NIPBL^c^ (WT or ATPase mutant) and dsDNA (5.19 kb) deposited onto a mica surface (+ 2.5 mM ATP). Both WT and ATPase mutant cohesin^SA1^-NIPBL^c^ complexes were randomly distributed on internal sites along dsDNA, with the WT complex displaying preferential binding to DNA ends (**Figure S1a&1b**). AFM images revealed different DNA binding modes by WT cohesin^SA1^-NIPBL^c^, as seen previously for condensin (33). Cohesin^SA1^-NIPBL^c^ molecules bound to DNA through the arm-hinge (**Figure 1d**, 30% ± 4.1%), the globular domain (**Figure 1e**, 53% ± 2.1%), both the arm-hinge and globular domains (**Figure 1f**, 15% ± 2.2%), or the foot (**Figure 1g**, 1.9% ± 0.2%, and **Figure 1h**). Furthermore, we observed similar DNA binding modes by the ATPase-deficient cohesin^SA1^-NIPBL^c^ EQ mutant in AFM images (+ATP, **Figure S1c-g**). These results, taken together, suggest that ATP hydrolysis is not needed for cohesin^SA1^-NIPBL^c^ loading onto DNA.

To further study how cohesin-NIPBL dynamically loads onto DNA and initiates a DNA loop, we applied HS-AFM imaging of WT and ATPase mutant cohesin^SA1^-NIPBL^c^ with dsDNA in liquids. Building on our previous success of using AFM imaging in liquids to study protein-DNA complexes (44,45), we developed robust sample deposition conditions on a 1-(3-Aminopropyl)silatrane treated mica (APS-mica) surface (41). We deposited WT cohesin^SA1^-NIPBL^c^ (30 nM) with DNA (3 nM, 5.19 kb) onto an APS-mica surface after 16-fold dilution and scanned the sample in a buffer containing ATP (+ 4 mM ATP) using either a Cypher VRS or JPK NanoWizard AFM at a scan rate of 0.4-2.3 frames/s. Importantly, under our sample deposition and imaging conditions, both proteins and DNA were mobile on the APS-mica surface. In time-lapse HS-AFM images, cohesin^SA1^-NIPBL^c^ displayed similar conformations, including I-shape, closed-ring, and folded arms with some showing protruded feet (gray arrows, **Figure 2a**), as observed in the static images collected in air (**Figure 1**). Analysis of individual cohesin^SA1^-NIPBL^c^ molecules interacting with DNA, recorded in real-time, revealed that cohesin^SA1^-NIPBL^c^ was highly dynamic in the presence of DNA (**Figure 2b**). Figure 2b shows one example of a monomeric WT cohesin^SA1^-NIPBL^c^ molecule with a bent elbow extending its arm-hinge domain to capture DNA in proximity (red arrow, compare **Figures 2b I, II**, and **III**). Both arms from this cohesin^SA1^-NIPBL^c^ molecule attempted to capture the DNA nearby (**Video S1**). Interestingly, a foot was also visible on this cohesin^SA1^-NIPBL^c^ molecule (gray arrow, **Figure 2b**), which transiently interacted with the DNA. In HS-AFM images, approximately 60% of cohesin ^SA1^-NIPBL^c^ molecules (N=50) showed either one- or two-foot structures.

**Figure 2.**
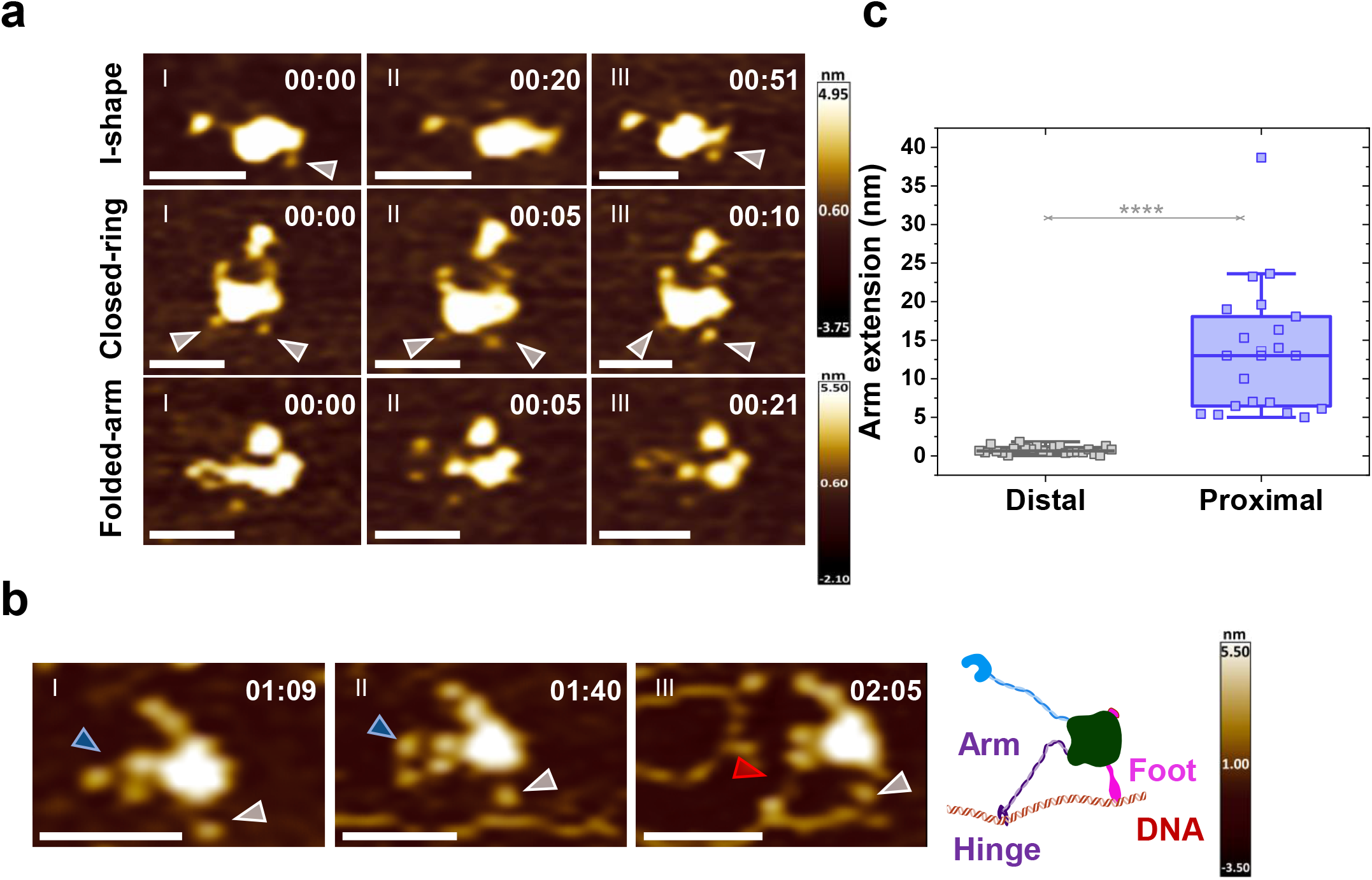
Real-time HS-AFM imaging in liquids reveals that WT cohesin-NIPBL captures DNA through the extension of the arm-hinge domain. (**a**) Time-lapse HS-AFM images showing diverse conformations of WT cohesin^SA1^-NIPBL^c^ in liquids (+ 4 mM ATP). (**b**) DNA capture by the extension of the arm-hinge domain on WT cohesin^SA1^-NIPBL^c^ (+ 4 mM ATP). DNA substrate: 5.19 kb. XY scale bar = 50 nm. See also **Video S1**. Blue arrow: arm; red arrow: arm extension; gray arrow: foot. Time: min:s. (**c**) Box-plot of arm extension lengths for WT cohesin^SA1^-NIPBL^c^ at a distal (> 500 nm distance, N=24 events) or proximal (< 50 nm distance, N=21 events) location from the DNA. Total: 3 experiments. 0.4-2.3 frame/s. Error bars: SD.

To investigate whether the presence of DNA drives arm extension, we further analyzed the change in arm lengths for WT cohesin^SA1^-NIPBL^c^ when the protein complex was either close (< 50 nm distance) or far (> 500 nm distance) from DNA. Strikingly, the arm-hinge extended significantly (p<2.5e-9) longer for cohesin^SA1^-NIPBL^c^ proximal to the DNA (N_proximal_=21, 13.6 nm ± 8.4 nm) compared to protein complexes distal to the DNA (N_distal_=24, 0.76 nm ± 0.5 nm, **Figure 2c**). In summary, HS-AFM imaging revealed unique protein conformational changes associated with DNA binding by cohesin-NIPBL.

Furthermore, we observed sequential events showing DNA being captured by the arm-hinge domain, followed by transferring of DNA to the globular domain and initiation of a DNA loop by WT cohesin^SA1^-NIPBL^c^ (**Figure 3**). This is exemplified, for instance, in Figure 3a, which shows that, after the DNA was captured by the arm of a cohesin^SA1^-NIPBL^c^ monomer (**Figure 3a I**), the arm swung backward (blue arrow, **Figure 3a II**), presumably bringing the DNA closer to the globular domain. Further, conformational changes of cohesin^SA1^-NIPBL^c^ led to the transfer of the DNA to the globular domain (compare the DNA location on cohesin^SA1^-NIPBL^c^ in **Figures 3a I** and **IV**). The second example in Figure 3b shows a cohesin^SA1^-NIPBL^c^ monomer with a closed-ring configuration that was initially proximal to the DNA (**Figure 3b I** and **Video S2**). The DNA was bent at an angle while being captured by the arm-hinge domain (**Figure 3b I**), then transferred to the globular domain (**Figure 3b II**). During the time interval when DNA was bound to the globular domain, the arm-hinge domains were open and extended out, trying to capture the nearby DNA at the top (red arrow, **Figure 3b III**) or the right (red arrow, **Figure 3b IV**). Notably, two feet were visible in some frames (gray arrows, **Figure 3b V, VI**, and **VII**), which transiently interacted with DNA (**Figure 3b VI**). Finally, a DNA loop was initiated after transient DNA binding by the foot and the capture of the nearby DNA segment by its arm-hinge domain (**Figure 3b VII**).

**Figure 3.**
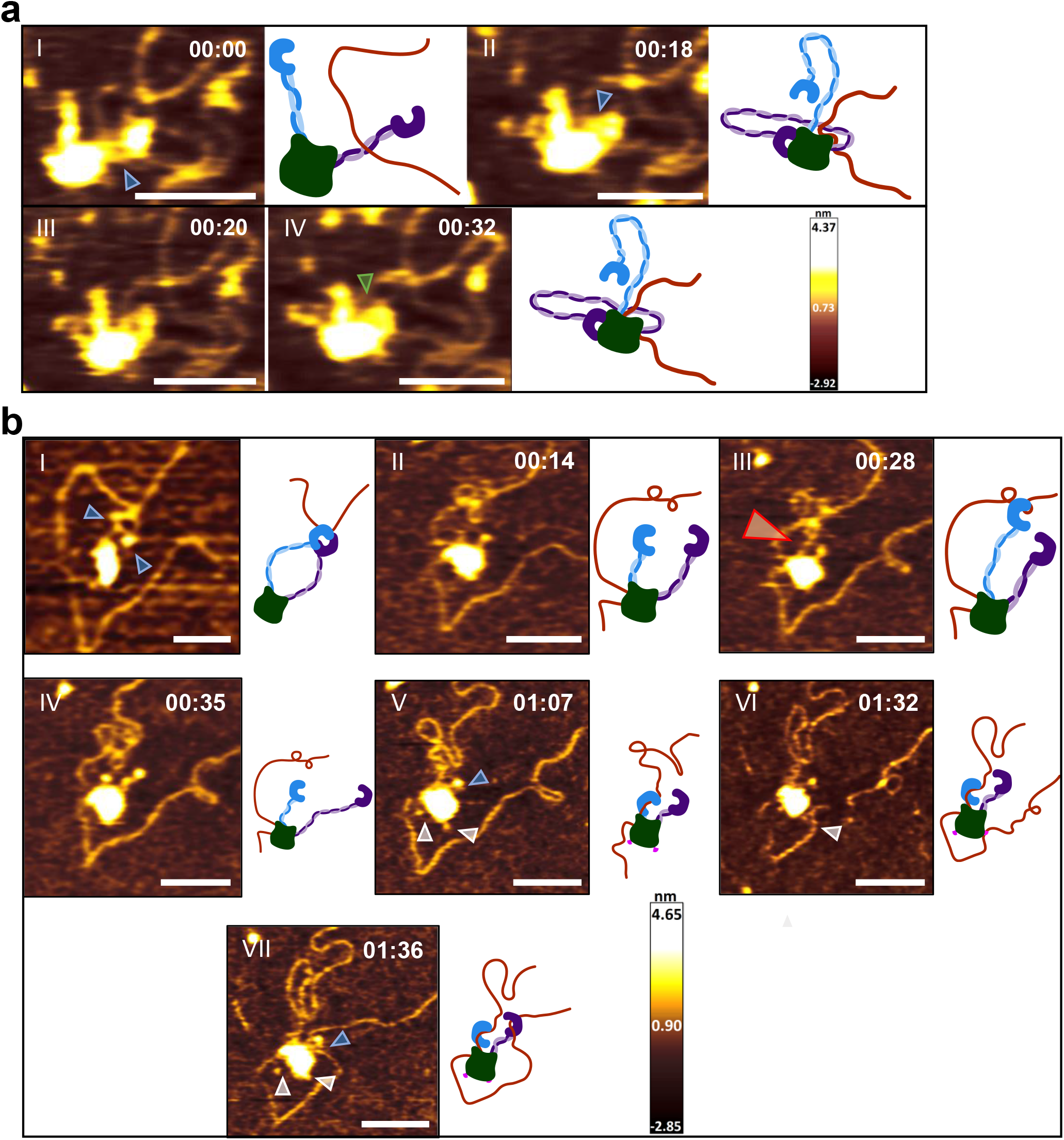
HS-AFM imaging in liquids shows sequential DNA binding events and the initiation of a DNA loop by WT cohesin-NIPBL. (**a** and **b**) Time-lapse HS-AFM images (left panels) and models (right panels) showing initial DNA capture by the arm-hinge domain, transfer of DNA binding to the globular domain, and (**b**) initiation of a DNA loop by the arm-hinge domain on WT cohesin^SA1^-NIPBL^c^ (+ 4 mM ATP). For panel 3b, also see **Video S2**. Panel bI is from an earlier time-lapse series of the same molecule. DNA substrate: 5.19 kb. XY scale bar = 50 nm. Blue arrow: arm; red arrow: arm extension; green arrow: binding at the globular domain; gray arrow: foot. Time: min:s.

While the cohesin^SA1^-NIPBL^c^ EQ ATPase mutant is expected to retain nucleotide-binding activity, it displays minimal ATPase catalytic activity in the presence of DNA and NIPBL^c^ (22). If initial DNA capture by the cohesin arm-hinge domain depends on ATPase hydrolysis, the cohesin^SA1^-NIPBL^c^ EQ ATPase mutant would be defective in arm extension. However, the cohesin^SA1^-NIPBL^c^ ATPase mutant displayed initial DNA search and capture processes similar to WT cohesin^SA1^-NIPBL^c^ (**Figure 4**). Figure 4a shows an example of a cohesin^SA1^-NIPBL^c^ ATPase mutant monomer displaying dynamic conformational changes before binding to DNA, including I-shape, closed-ring, and open-arms with bent elbows (**Figure 4a** and **Video S3**) (25). Like the WT protein, cohesin^SA1^-NIPBL^c^ ATPase mutant captured DNA through dramatic conformational changes and extension of the arm-hinge domain (red arrows in **Figures 4b&c** and **Videos S4**&**S5**). The average length of arm extension for the cohesin^SA1^-NIPBL^c^ ATPase mutant proximal to the DNA was measured to be 13.2 (± 8.6) nm (**Figure 4d**), comparable to the WT cohesin complex (**Figure 2c**). In some cases, the initial capture of DNA by the arm-hinge domain led to subsequent DNA loop formation (**Figure 4c III** and **Video S5**). Interestingly, HS-AFM imaging revealed diffusion (walking) of the cohesin^SA1^-NIPBL^c^ ATPase mutant on DNA using short protrusions (**Figure S2** and **Videos S3**&**S4**).

**Figure 4.**
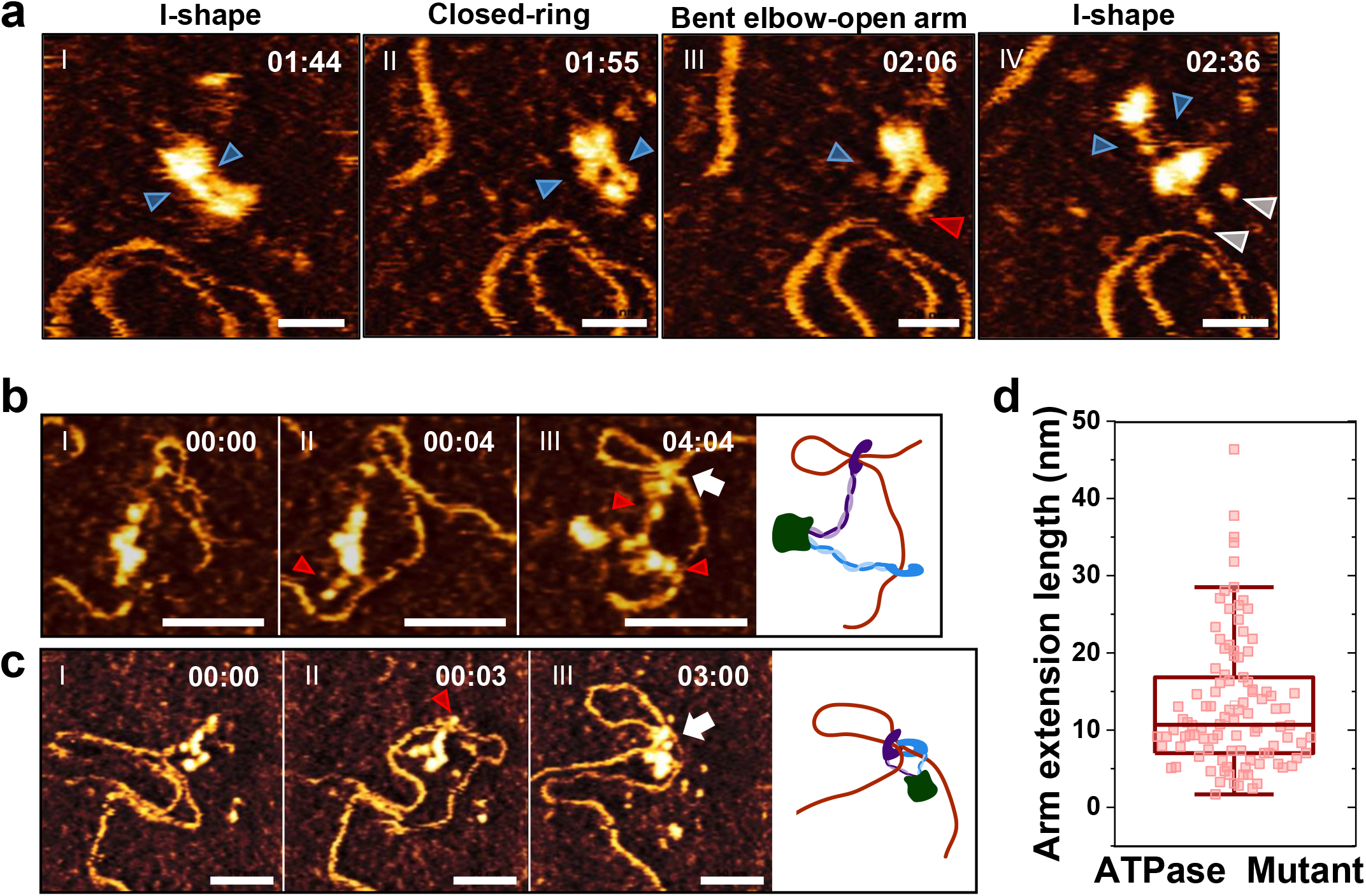
HS-AFM imaging in liquids demonstrates that the cohesin-NIPBL ATPase mutant captures DNA and initiates a DNA loop through the extension of the arm-hinge domain. (**a**) Conformational changes of cohesin^SA1^-NIPBL^c^ ATPase mutant (**Video S3**). (**b** and **c**) DNA capture through arm extension by the cohesin^SA1^-NIPBL^c^ ATPase mutant and initiation of a DNA loop (b: **Video S4**; c: **Video S5**). Blue arrow: arm; red arrow: arm extension; gray arrow: foot; white arrow: protein-mediated DNA loop. XY scale bar = 50 nm. Right panels in b and c: models. Time: min:s. Buffer contains 4 mM ATP. (**d**) Box plot showing arm extension lengths on the cohesin^SA1^-NIPBL^c^ ATPase mutant during DNA capture (13.2 nm ± 8.6 nm, N=102 events).

In summary, AFM imaging reveals that the SMC arm-hinge domain on cohesin-NIPBL is highly flexible and dynamic. Cohesin-NIPBL can capture DNA segments in proximity through the extension of the arm-hinge domain in an ATP hydrolysis-independent manner. The initial capture of DNA by the arm-hinge domain on cohesin-NIPBL may be followed by transferring of DNA to the globular domain and the initiation of a DNA loop through capturing the second DNA segment by the arm-hinge. Furthermore, feet protruding from the globular domain can transiently bind to DNA and facilitate DNA segment capture by the arm-hinge domain.

### ATP-independent and dependent cohesin-NIPBL mediated DNA looping and bending

HS-AFM imaging shows that both WT and ATPase mutant cohesin^SA1^-NIPBL^c^ can form DNA loops through diffusion capture of DNA segments in proximity (**Figures 3b** and **4c**). Next, we directly compared the DNA looping efficiency and loop structures mediated by WT and ATPase mutant cohesin^SA1^-NIPBL^c^. AFM images (collected in air) of WT (± ATP) and ATPase mutant (+ATP) cohesin^SA1^-NIPBL^c^ (30 nM) in the presence of dsDNA (5.19 kb, 6 nM) showed distinct protein-mediated DNA loops (**Figure 5a**). On incubating WT (-ATP) or ATPase mutant cohesin^SA1^-NIPBL^c^ (+ATP) with the linear dsDNA, 15.6% (±4.3%) and 18.1% (±0.3%) of dsDNA molecules, respectively, contained protein-mediated DNA loops (**Figure 5b**). For WT cohesin^SA1^-NIPBL^c^, the addition of ATP (+2.5 mM ATP) significantly increased (p<0.05) the population of DNA molecules with protein-mediated loops to 65.2% (±3.6%, **Figure 5b**).

**Figure 5.**
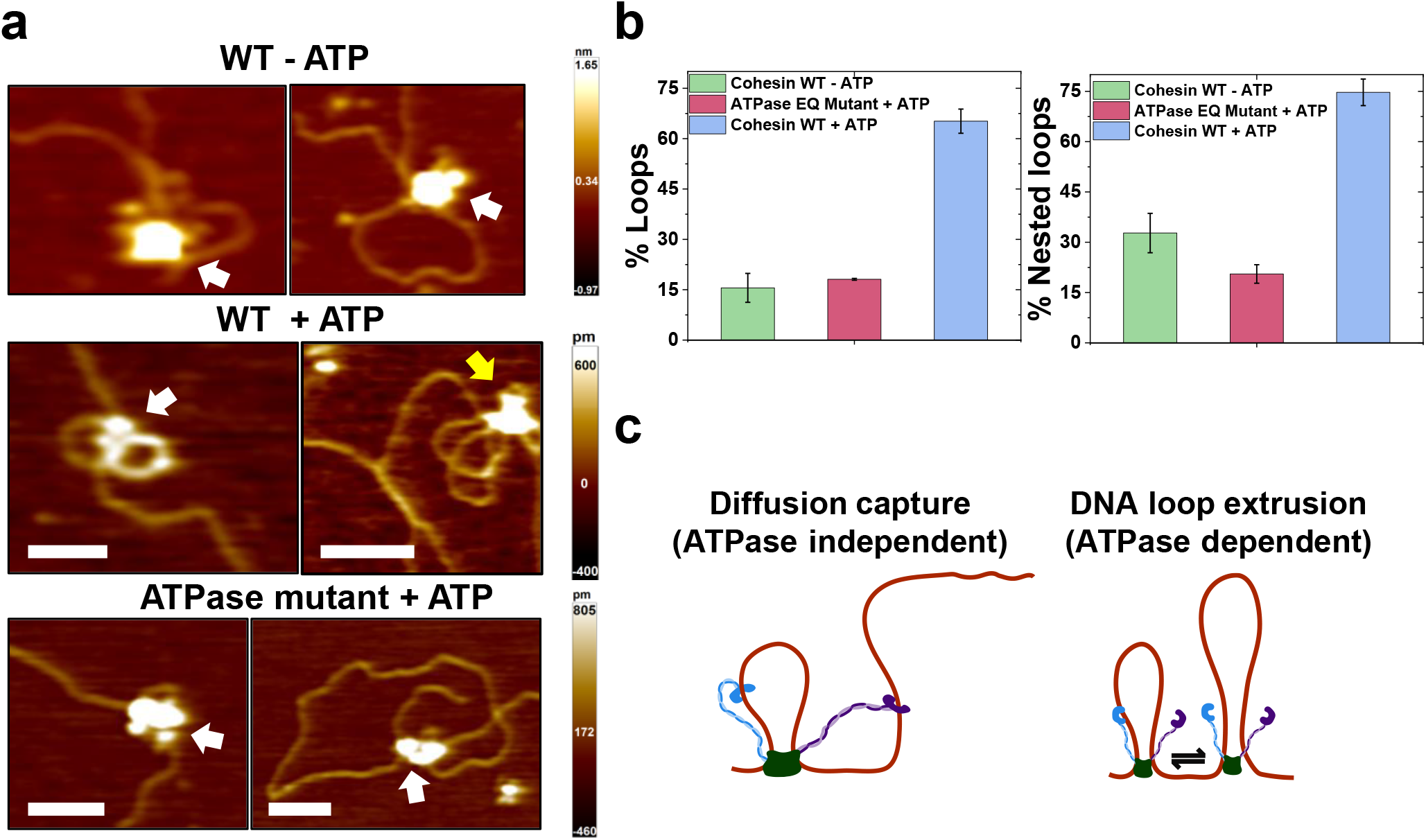
AFM imaging in air reveals cohesin-NIPBL mediated DNA loops. (**a**) Representative AFM images of DNA loops mediated by WT cohesin^SA1^-NIPBL^c^ in the absence (top) and presence of ATP (middle, 2.5 mM ATP), and the cohesin^SA1^-NIPBL^c^ ATPase mutant in the presence of ATP (bottom, 2.5 mM ATP) on linear DNA. Cohesin^SA1^-NIPBL^c^: 30 nM. DNA (5.19 kb): 6 nM. White arrow: single loop; Yellow arrow: nested loop. XY scale bar = 100 nm. (**b**) Quantification of the percentages of DNA molecules containing protein-mediated total DNA loops (left panel, N=155) and nested loops (right panel, N=119). Error bars: SD. 2 experiments for each condition. (**c**) A model representing mechanisms of ATPase-independent diffusion capture of an additional DNA segment (left panel) and ATPase-dependent DNA loop extrusion by cohesin-NIPBL that might lead to nested DNA loops (right panel).

Furthermore, AFM imaging revealed cohesin^SA1^-NIPBL^c^ mediated nested DNA loops (a loop within a loop, yellow arrows in **Figure 5a**). Nested DNA loops can be generated when cohesin-NIPBL at an existing DNA loop capture an additional DNA segment (ATPase-independent) or two separate cohesin-NIPBL molecules on the same DNA collide after DNA loop extrusion (ATPase-dependent, **Fig. 5c**) (46). The population of nested DNA loops observed for WT cohesin^SA1^-NIPBL^c^ in the presence of ATP (74.7%±4.0%) was significantly (p<0.05) greater than that observed for either WT cohesin^SA1^-NIPBL^c^ without ATP (32.8%±5.9%) or the ATPase mutant (20.6%±2.8%, **Figure 5b**). The nested loops formed by WT cohesin^SA1^-NIPBL without ATP or the ATPase mutant could be due to the capture of additional DNA segments at an existing protein-mediated DNA loop in an ATPase-independent manner (**Figure 5c**). These results collectively suggest that cohesin-NIPBL mediates DNA loops through two distinct mechanisms: ATPase-independent diffusion capture of DNA segments in proximity and ATPase-driven DNA loop extrusion (**Figure 5c**).

In addition to DNA loops, AFM imaging in air revealed cohesin^SA1^-NIPBL^c^-induced DNA bending (**Figure S3**). While DNA alone showed slight bending (27.4° ± 26.0°, +ATP) measured at randomly chosen positions, WT cohesin^SA1^-NIPBL^c^ in the absence of ATP (43.7° ± 20.5°) and cohesin^SA1^-NIPBL^c^ ATPase mutant (+ATP, 47.3° ± 41.0°) induced significantly (p<0.05) higher degrees of DNA bending (**Figure S3d**). Furthermore, compared to DNA binding by WT cohesin^SA1^-NIPBL^c^ without ATP, the presence of ATP further augmented (p<0.05) the DNA bending (57.2° ± 27.6°, **Figure S3d**). Additionally, we compared the DNA bending angles induced by either the globular or the hinge domain. The globular domain on WT and ATPase mutant cohesin^SA1^-NIPBL^c^ induced comparable DNA bending, which was significantly (p<0.05) higher than what was induced by the hinge domains (**Figure S3e**). In summary, these results from AFM imaging demonstrate that cohesin-NIPBL bends DNA independent of ATP hydrolysis at different DNA binding steps, which could facilitate DNA looping.

### HS-AFM imaging in liquids reveals DNA loop extrusion dynamics by cohesin-NIPBL

AFM imaging in air revealed that WT cohesin^SA1^-NIPBL^c^ in the presence of ATP induced a higher percentage of DNA loops than the WT protein complex without ATP or the ATPase mutant (**Figure 5**). This result is consistent with the notion that WT cohesin^SA1^-NIPBL^c^ is capable of DNA loop extrusion in an ATP hydrolysis-dependent manner. A recent magnetic tweezers study of condensin with a resolution of ∼10 nm revealed a broad distribution of DNA looping step sizes (47). We expected that real-time HS-AFM imaging of cohesin^SA1^-NIPBL^c^ with DNA (+ATP) would directly reveal DNA extrusion steps. Because DNA movement during imaging could contribute to slight DNA length fluctuations without DNA loop extrusion, we first carried out control experiments using HS-AFM imaging to measure DNA loop length changes for the cohesin^SA1^-NIPBL^c^ ATPase mutant (+ATP, **Figure S4**). The DNA loop lengths fluctuated slightly with small forward (increased, 1.2 nm ± 1.1 nm, N=60 events) and reverse (decreased, - 1.34 nm ± 1.0 nm, N=51 events) changes (per second). The total DNA loop lengths did not change significantly over time (N=7 DNA loops, **Figure S4**). In stark contrast, DNA loop lengths mediated by WT cohesin^SA1^-NIPBL^c^ (+ATP, N=18 DNA loops) showed forward and reverse step sizes, significantly higher than the background fluctuation observed for the ATPase mutant (**Figure 6, Figure S5**, and **Videos S6**&**S7**). The distribution of the DNA looping step size mediated by WT cohesin^SA1^-NIPBL^c^ greater than the background fluctuation (>5 nm) displayed forward steps at 13.2 nm (± 16.1 nm, N=115) and reverse steps at -12.0 nm (± 9.8 nm, N=107, **Figure 6d**). Collectively, HS-AFM imaging demonstrates active DNA loop extrusion by cohesin-NIPBL with distinct DNA loop extrusion step sizes.

**Figure 6.**
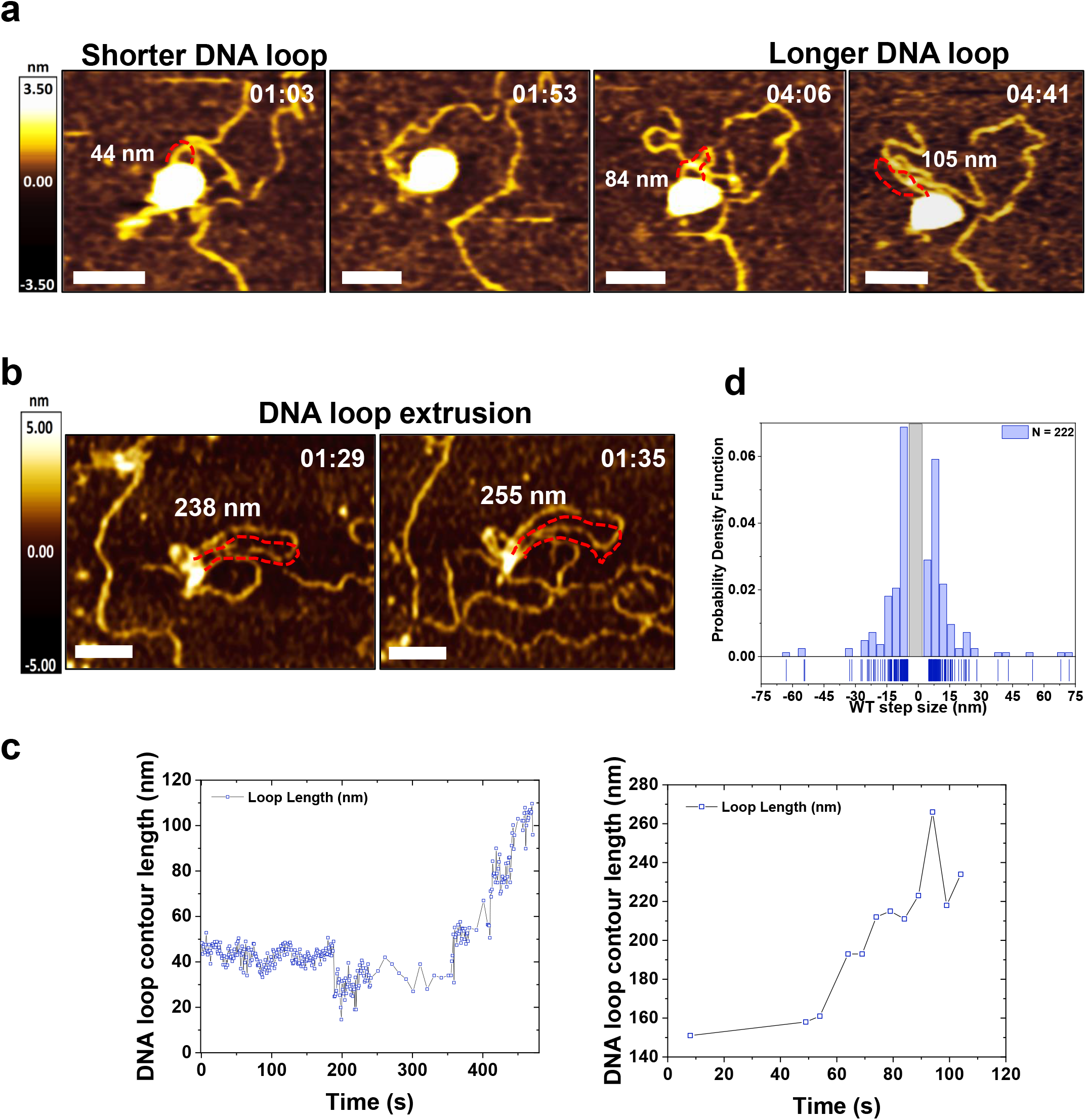
DNA loop extrusion revealed by HS-AFM imaging of WT cohesin-NIPBL-DNA complexes in the presence of ATP. (**a** and **b**) Time-lapse AFM images showing DNA loop length changes mediated by WT cohesin^SA1^**-**NIPBL^c^ on a linear dsDNA (5.19 kb) in a buffer containing 4 mM ATP. Time: min:s. XY scale bar = 50 nm. Dotted red lines mark the DNA loops and the numbers in nm indicate DNA loop lengths. (**c**) The DNA loop contour lengths over time measured for the time-lapse image series shown in panels a (left panel, **Video S6**) and b (right panel, **Video S7**). (**d**) The forward (13.2 nm ± 16.1 nm, N=115) and reverse (−12.0 nm ± 9.8 nm, N=107) DNA loop extrusion step size (per second) measured based on frame-to-frame loop length changes in HS-AFM images (15 loops, 40 extrusion events, and four technical repeats). Background fluctuation of DNA length (<5 nm) based on the measurement for the cohesin^SA1^-NIPBL^c^ ATPase mutant (**Figure S4**) was excluded (gray rectangle in panel d). 1-2.3 frames/s.

### DNA binding by the C-terminal domain of cohesin SA2

A recent study identified three dsDNA binding patches on SA1, including Patch 1 (K92, K95, K172, and K173), 2 (K555, K558, and R564), and 3 (K969, R971, K1013, and R1016) (25). However, DNA binding by the C-terminal domains of SA1/SA2 and NIPBL, which are disordered in the cryo-EM structure of cohesin-NIPBL (42), has not been investigated. SA1 and SA2 are highly similar, with approximately 70% sequence identity (48). To further establish DNA binding domains on SA1/SA2, we purified WT full-length SA2 and SA2 fragments, including the N-terminal (1-301 AAs or 1-450 AAs), middle (451-1051 AAs), and C-terminal (1052-1231 AAs) domains (**Figure S6**) (37). Fluorescence anisotropy measurements using a fluorescently labeled dsDNA substrate (45 bp) revealed that SA2 contains extensive DNA binding surfaces. All three (N-terminal, middle, and C-terminal) domains bind to dsDNA, albeit with the highest binding affinities contributed by the N-terminal domain (1-302 AAs: K_d_=359 nM; 1-450 AAs: K_d_=358 nM). Notably, the middle (451-1051 AAs: K_d_=1134 nM) and C-terminal domain of SA2 (1052-1231 AAs: K_d_=5962 nM) bind to dsDNA weakly. Consistent with these results, we showed previously that deletion of the C-terminal domain of SA2 (SA2 1-1051 AAs) reduces its binding affinity for dsDNA (39). Thus, these results from fluorescence anisotropy suggest that the C-terminal domain of SA1/SA2 has the potential to bind DNA, and likely contributes to one of the feet protruding from the globular domain observed in AFM images. Indeed, the C-terminal domain of SA1/SA2 can easily get cleaved during protein purification, suggesting that this domain has an extended structure, consistent with it appearing as the foot-like feature observed in AFM images.

## DISCUSSION

### Extension of cohesin arm-hinge mediates dynamic DNA search and initiation of DNA loops by cohesin-NIPBL

While recent single-molecule fluorescence studies demonstrated DNA loop extrusion by cohesin-NIPBL, the mechanisms of DNA capture and DNA loop initiation by cohesin-NIPBL are still under intense debate (49-51). Here, we used traditional AFM and HS-AFM imaging to directly observe these initial DNA interactions by cohesin-NIPBL. It is worth noting that HS-AFM imaging in liquids relies on an intricate balance to keep protein and DNA molecules partially anchored onto a surface while still being mobile. What this entails is that for each protein complex, we might not be able to observe its full range of motion and the complete process from DNA loading to DNA loop extrusion. However, HS-AFM provides a unique window into sequential events during DNA binding and protein conformational changes associated with DNA binding (**Figure 7**). The crystal structure of the SMC1-SMC3 hinge heterodimer contains a short ssDNA bound to the outer surface of the SMC1 hinge, suggesting its role in DNA binding (42). In this study, HS-AFM imaging in liquids reveals dynamic search and DNA capture by the cohesin arm-hinge domain. The cohesin^SA1^-NIPBL^c^ ATPase mutant displays the same DNA capture process through the arm-hinge domain, suggesting that it is partly driven by Brownian motion instead of ATP hydrolysis. Previously, we solved two structures of the SMC1-SMC3 hinge heterodimer that adopt different open conformations, suggesting that the interface between human SMC1-SMC3 hinges is highly dynamic (42). Strikingly, HS-AFM imaging shows that the arm-hinge of WT and ATPase mutant cohesin^SA1^-NIPBL^c^ is extended ∼14 nm to capture DNA in proximity. The SMC hinge domains contain positively charged patches (25). Likely, the electrostatic interaction between the hinge domain and negatively charged DNA backbone drives DNA binding at the hinge domain. This model is consistent with previous findings that mutations at three conserved lysine residues on the lumen of the yeast cohesin abolished the loading of cohesin onto the chromatin (52). Based on these observations, the Nasmyth group suggested that the positive charges normally hidden inside the SMC hinge’s lumen are transiently exposed to DNA through significant conformational changes of the arm-hinge domain (52). Capturing the second DNA segment by the arm-hinge of cohesin could contribute to the bridging of sister chromatids and cohesion. Consistent with this notion, mutations in the yeast SMC1 and SMC3 hinge domains that neutralize a positively charged channel led to sister chromatin cohesion defects (53). It is worth noting that the Debye screening length around dsDNA is ∼1 nm at the ionic strength used for HS-AFM imaging. Since both protein and DNA molecules were mobile during AFM imaging, the precise distance between cohesin and DNA that activates arm extension could be significantly shorter than what we can measure based on AFM images.

**Figure 7.**
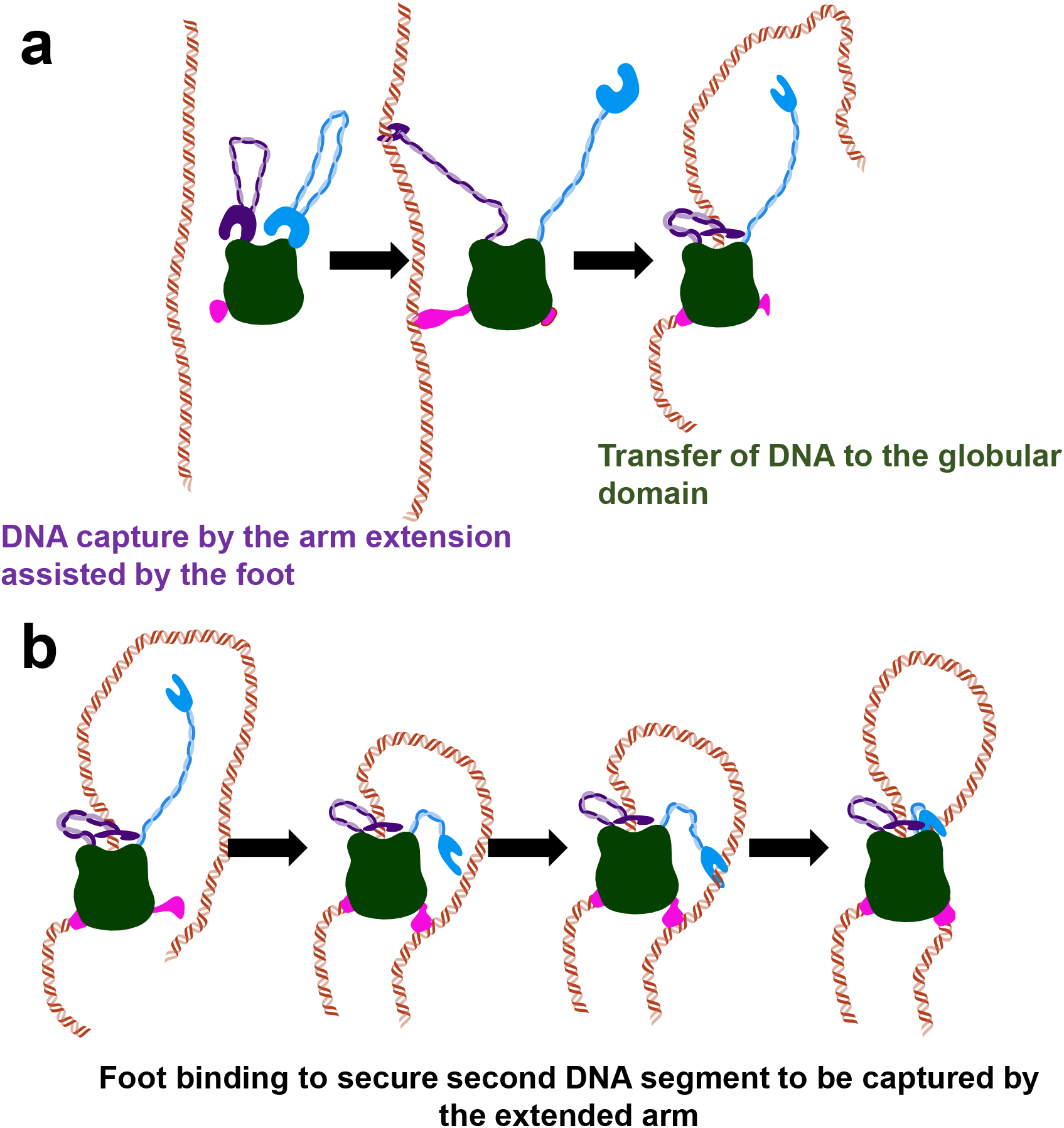
Multi-step DNA binding and loop initiation model for cohesin-NIPBL. (**a**) DNA capture by arm extension followed by transferring of DNA to the globular domain in an ATPase-independent manner. (**b**) Transient DNA binding by the foot brings DNA closer to cohesin-NIPBL, facilitating DNA capture by the arm-hinge domain.

Consistent with previous studies (25), in our AFM images, SMC1 and SMC3 heads, RAD21, SA1, and NIPBL^c^ collectively show up as a globular particle (globular domain). DNA binding surfaces on these subunits have been revealed by cryo-EM structures of cohesin^SA1^-NIPBL^c^ and DNA binding assays in conjunction with mutagenesis (25,42). Upon initial DNA binding through the SMC arm-hinge domain, DNA is transferred to the globular domain (**Figure 7a**), for which our recent cryo-EM structure of cohesin^SA1^-NIPBL^c^ provides additional detail on DNA binding (42). Specifically, this structure showed that cohesin^SA1^-NIPBL^c^ binds DNA at the top of the engaged SMC1-SMC3 heads with NIPBL and SA1 wrapping around DNA, creating a central channel (42). It was suggested that ATP binding opens the head gate to complete the DNA entry, and head engagement leads to a DNA “gripping/clamping” state (51). Results from HS-AFM imaging from this study do not contradict this model. Instead, observations from this study support a comprehensive model in which DNA search and transient DNA binding by the arm-hinge precede the DNA “gripping/clamping” state at the globular domain.

### Cohesin-NIPBL feet and function

Unexpectedly, in addition to arms, some cohesin^SA1^-NIPBL^c^ molecules display short protrusions (feet) from the globular domain. When these feet are in the proximity of DNA, they transiently bind to DNA, holding DNA closer to cohesin^SA1^-NIPBL^c^, thereby facilitating the capture of the DNA by the arm-hinge domain (**Figure 7**). We also observed a random walk of cohesin^SA1^-NIPBL^c^ on DNA through short protrusions (likely feet), driven by thermal energy. These feet are likely the C-terminal domains of SA1 and NIPBL (∼200 AAs), which are unstructured in the cohesin^SA1^-NIPBL^c^ cryo-EM structures (42). This notion is supported by the observation of DNA binding by the C-terminal domain of SA2 (1051-1231 AAs). Furthermore, sequence alignment shows that the C-terminal domain of NIPBL contains numerous conserved positively charged residues (**Figure S7**).

### Two distinct and collaborative DNA looping mechanisms by cohesin-NIPBL

AFM imaging of cohesin-NIPBL complexes from this study demonstrates that cohesin-NIPBL promotes DNA looping through two distinct mechanisms. WT cohesin^SA1^-NIPBL^c^ without ATP and the ATPase mutant are both capable of capturing DNA loops. These results show that cohesin-NIPBL can sequentially capture two DNA segments in proximity through Brownian motion (diffusion capture) independent of ATP binding and hydrolysis (**Figure 7b**) (54). Secondly, WT cohesin^SA1^-NIPBL^c^ in the presence of ATP further increases the percentage of DNA molecules displaying loops and nested loops, likely through ATPase-dependent DNA loop extrusion (**Figure 5c**). Molecular dynamics simulations showed that a combination of diffusion capture and loop extrusion recapitulates condensin-dependent mitotic chromatin contact changes (54). These two DNA looping pathways could also function collaboratively through ATPase-dependent DNA loop extrusion following diffusion capture of DNA by cohesin-NIPBL.

### DNA loop extrusion dynamics by cohesin-NIPBL

Despite recent experimental demonstrations of DNA loop extrusion by cohesin and condensin *in vitro* and *in cellulo* (22,23,25,46,55-58), we have not reached a consensus regarding the molecular mechanism of DNA loop extrusion (36). Several competing DNA loop extrusion models have been proposed, including the tethered-inchworm (34), DNA-segment-capture (35), hold-and-feed (59), scrunching (33), swing and clamp (25), and Brownian ratchet models (36). HS-AFM imaging in this study demonstrates that once DNA is bound to the globular domain in the DNA “gripping/clamping” state (25), the SMC arm-hinge domain of both WT and ATPase mutant cohesin^SA1^-NIPBL^c^ is free to search the next DNA fragment through arm extension, bending (“swinging”) towards the head domain, leading to initiation of a DNA loop. These results show that it is not the ATP hydrolysis or power stroke that drives arm extension and capture of the DNA segment. Furthermore, consistent with our findings, artificially induced stable bending of the hinge towards the head inhibits DNA loop extrusion by cohesin-NIPBL (25). The conformational change of cohesin-NIPBL driving DNA loop growth is still hotly debated. The Brownian ratchet model postulates that loop growth depends on the stochastic Brownian motion of the Scc3-hinge domain, followed by DNA slipping along the Scc2-head domain (26). The swing and clamp model posits that DNA translocation and loop growth is through the synchronization of the head-disengagement/engagement driven by the ATPase cycle and arm-hinge swing/DNA clamping (25). While HS-AFM imaging does not provide detail on relative movements of the SMC head domains, SA1, and NIPBL during DNA loop extrusion, it shows DNA loop extrusion with cohesin^SA1^-NIPBL^c^ partially anchored to a surface. This result suggests a mechanism that relies on cohesin-NIPBL switching between DNA gripping and slipping states where DNA can slide across the cohesin-NIPBL globular domain leading to DNA loop growth.

It is known that tension on DNA reduces the DNA loop extrusion step size (47). Consistent with this notion, since DNA was partially anchored onto a surface that likely generates tension, we observed “bursts” of DNA loop extrusion events when the tension on DNA was favorable in HS-AFM imaging. On the other hand, with nanometer spatial resolution, HS-AFM enabled observations of DNA looping step sizes. Although the DNA looping step size measured from HS-AFM images (∼13 nm or 42 bp) is considerably lower than what is estimated by combining the loop extrusion speed (∼0.5-1 kb/s) and ATPase rate (2 ATP/s) (22,23), it is similar to the step size of condensin under DNA stretching forces from 1 to 0.2 pN (∼20-40 nm)(47). In addition, HS-AFM imaging shows both forward and reserve steps, suggesting that cohesin-NIPBL can switch DNA strands during DNA loop extrusion. It is highly likely that surface anchoring of DNA and cohesin-NIPBL increases the frequency of strand switching and DNA loop extrusion pausing (40,60-67).

In summary, HS-AFM imaging reveals dynamic conformational changes on cohesin-NIPBL that drive DNA loading and loop formation. This study uncovers critical missing links in our understanding of cohesin-NIPBL DNA binding and DNA loop extrusion (68,69).

## DATA AVAILABILITY

The data that support this study are available from the corresponding authors upon reasonable request.

## ACKNOWLEDGEMENTS

We would like to thank Ryan Fuierer and Keith Jones at Asylum Research and Dorothy Erie at the University of North Carolina at Chapel Hill for access to HS-AFM and technical support.

## AUTHOR CONTRIBUTIONS

P.K. designed and carried out the AFM experiments, analyzed the data, and wrote the paper. Z.S. purified the cohesin proteins and performed the sequence alignment. X.L. performed the fluorescence anisotropy experiments and analyzed the data. H.Z., I.J.F., Y.J.T., H.Y., and H.W. designed the experiments and wrote the paper.

## Supporting Information

**Video S1. HS-AFM video showing dynamics of the WT cohesin**^**SA1**^**-NIPBL**^**c**^ **and extension of the arm-hinge domain to capture a DNA segment in proximity (4 mM ATP)**. Related to Figure 2b.

**Video S2. HS-AFM video showing that a DNA-bound WT cohesin**^**SA1**^**-NIPBL**^**c**^ **complex captures a DNA segment in proximity through the arm-hinge domain and initiates a DNA loop (4 mM ATP)**. Related to Figure 3b.

**Video S3. HS-AFM video showing dynamics of cohesin**^**SA1**^**-NIPBL**^**c**^ **ATPase mutant and diffusion on DNA through short protrusions (4 mM ATP)**. Related to Figures 4a and S2a.

**Video S4. HS-AFM video demonstrating diffusion on DNA through short protrusions and arm extension by the cohesin**^**SA1**^**-NIPBL**^**c**^ **ATPase mutant (4 mM ATP)**. Related to Figure 4b (arm extension) and Figure S2b (walking on DNA).

**Video S5. HS-AFM video showing DNA capture by the cohesin**^**SA1**^**-NIPBL**^**c**^ **ATPase mutant through arm extension and initiation of a DNA loop (4 mM ATP)**. Related to Figure 4c.

**Video S6. HS-AFM video showing DNA loop extrusion by WT cohesin**^**SA1**^**-NIPBL**^**c**^ **(4 mM ATP)**. Related to Figure 6a.

**Video S7. HS-AFM video showing DNA loop extrusion by WT cohesin**^**SA1**^**-NIPBL**^**c**^ **(4 mM ATP)**. Related to Figure 6b.

**Figure S1.**
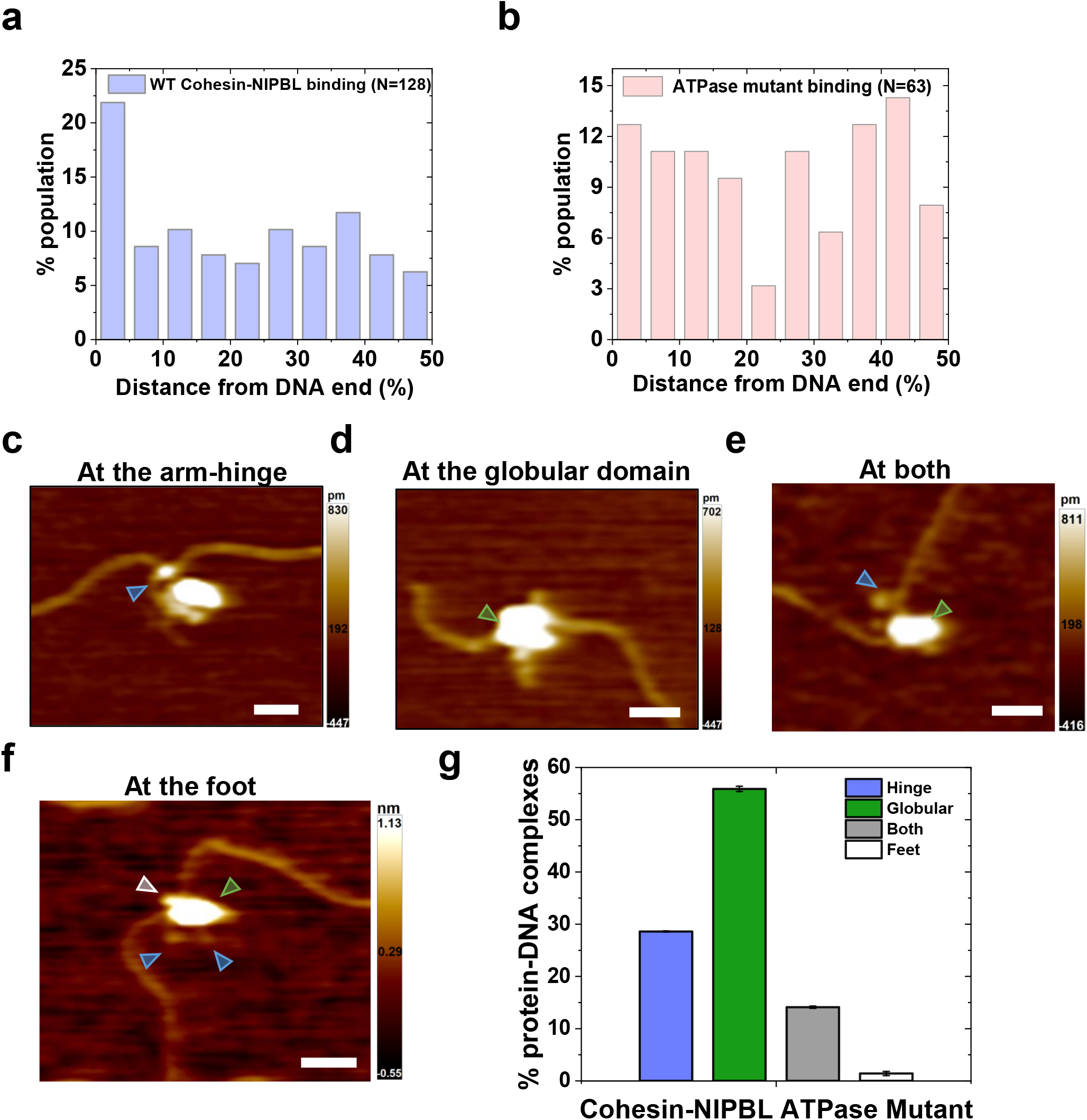
DNA binding position distributions of WT and ATPase mutant cohesin^SA1^-NIPBL^c^ on dsDNA and DNA binding modes by ATPase mutant cohesin^SA1^-NIPBL^c^. (**a** and **b)** Position distributions of WT (**a**) and ATPase mutant (**b**) cohesin^SA1^-NIPBL^c^ on dsDNA (5.19 kb). (**c** to **f**) Example AFM images of cohesin^SA1^-NIPBL^c^ ATPase mutant binding DNA at the arm-hinge (**c**), globular domain (**d**), both the arm-hinge and globular domains (**e**), and the foot (**f**). XY scale bar = 50 nm. Blue arrow: arm; green arrow: globular domain; gray arrow: foot. (**g**) Percentages of ATPase mutant cohesin^SA1^-NIPBL^c^-DNA complexes with the arm-hinge (28.6% ± 0.1%), globular (55.9% ± 0.5%), both arm-hinge and globular domains (14.1% ± 0.2%), and foot (1.4% ± 0.4%) binding to DNA. N=157 protein-DNA complexes. Error bars: SD. Two independent experiments in the presence of 2.5 mM ATP.

**Figure S2.**
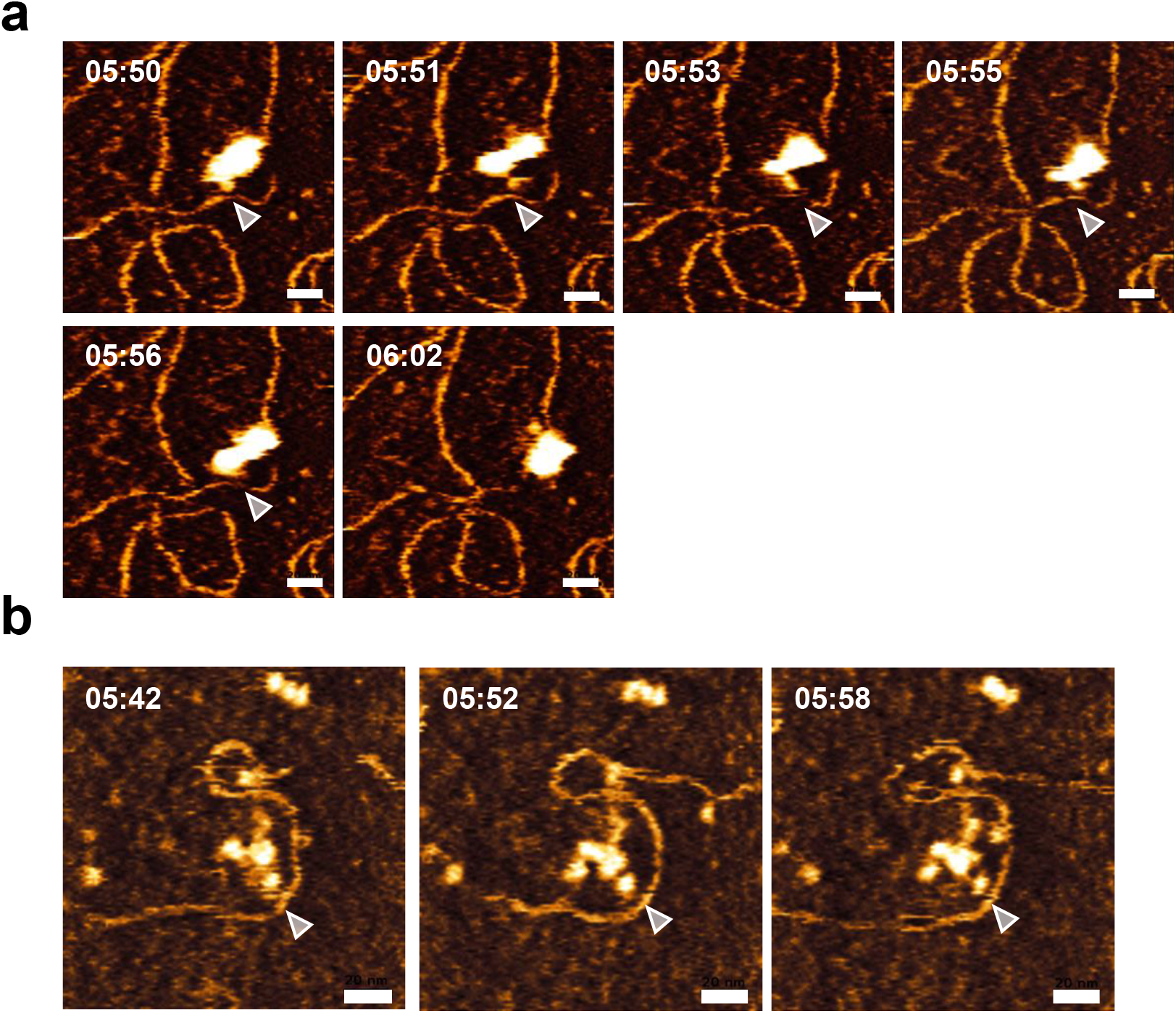
Time-lapse HS-AFM images of cohesin^SA1^-NIPBL^c^ ATPase mutant walking (diffusion) on DNA through short protrusions. XY scale bar = 20 nm. Also see **Video S3** (panel a) and **Video S4** (panel b). Panels a and b observation times are in continuation of Figure 4a and 4b, respectively, for the same molecules. gray arrow: foot. Time: min:s.

**Figure S3.**
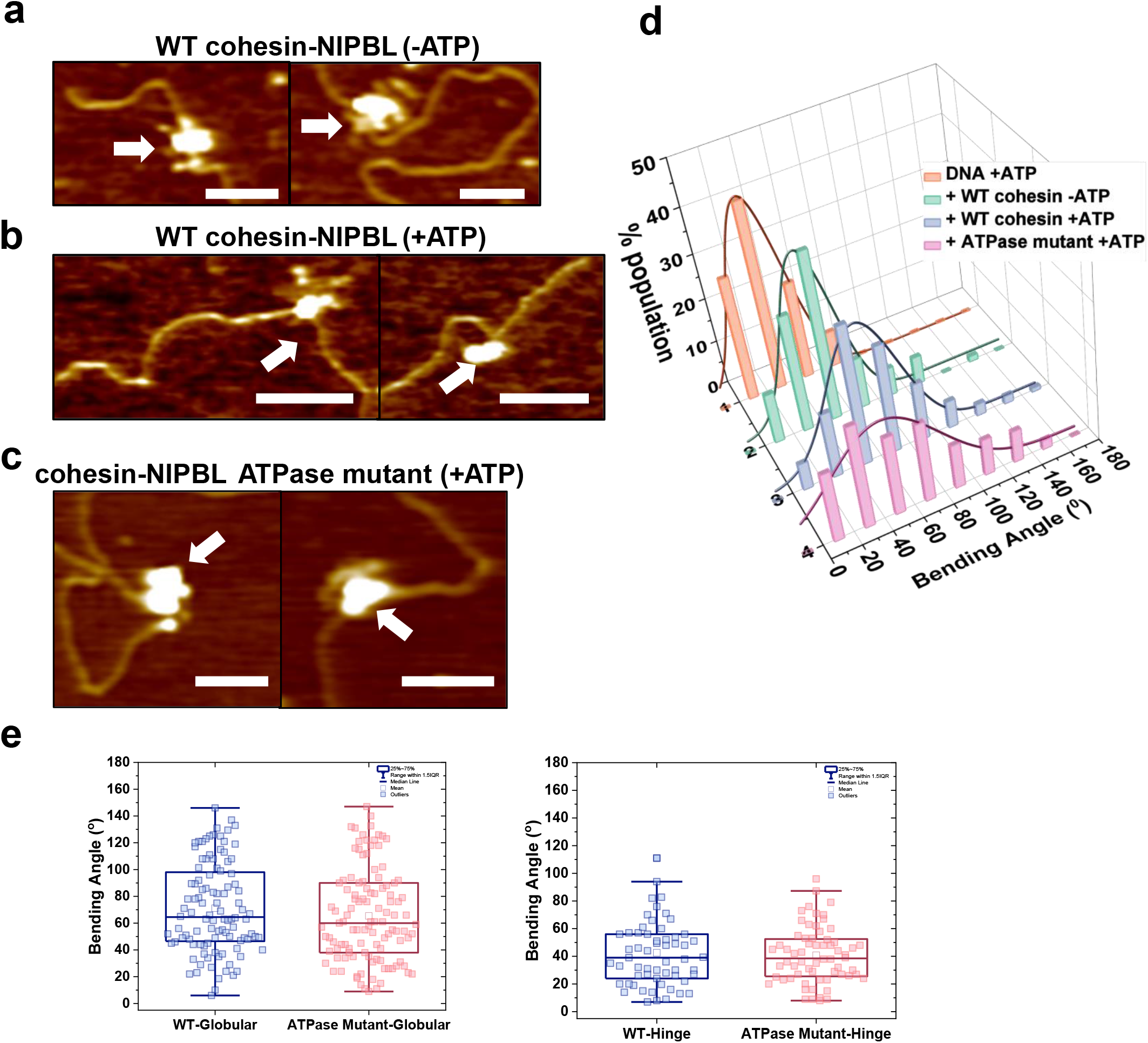
Cohesin^SA1^-NIPBL^c^ binding induces DNA bending. (**a** to **c**) Representative AFM images of WT cohesin^SA1^-NIPBL^c^ in the absence (**a**), or presence of ATP (+2.5 mM, **b**), and cohesin^SA1^-NIPBL^c^ ATPase mutant (+2.5 mM ATP, **c**) with dsDNA (5.19 kb). White arrows point to protein-DNA complexes. XY scale bar = 100 nm. (**d**) Histograms of the DNA bending angles induced by the WT and ATPase mutant of cohesin^SA1^-NIPBL^c^ on dsDNA. The solid lines are Gaussian fits (R^2^> 0.8) to the data with peaks centered at the DNA bending angle of 27.5° (± 26°) for DNA_+ATP_ (N=149), 43.7° (± 20.5°) for DNA_WT-ATP_ (N=80), 57.2° (± 27.6°) for DNA_WT+ATP_ (N=116), and 47.3° (± 41.0°) for DNA_ATPase mutant+ATP_ (N=298). (**e**) DNA bending angles induced by the globular domain of the WT (71.2° ± 33.6°, N=104) and ATPase mutant (65.4° ± 34.6°, N=111), and the hinge domain of WT (42.6° ± 24.4°, N=59) and ATPase mutant (40.7° ± 20.3°, N=104) cohesin^SA1^-NIPBL^c^.

**Figure S4.**
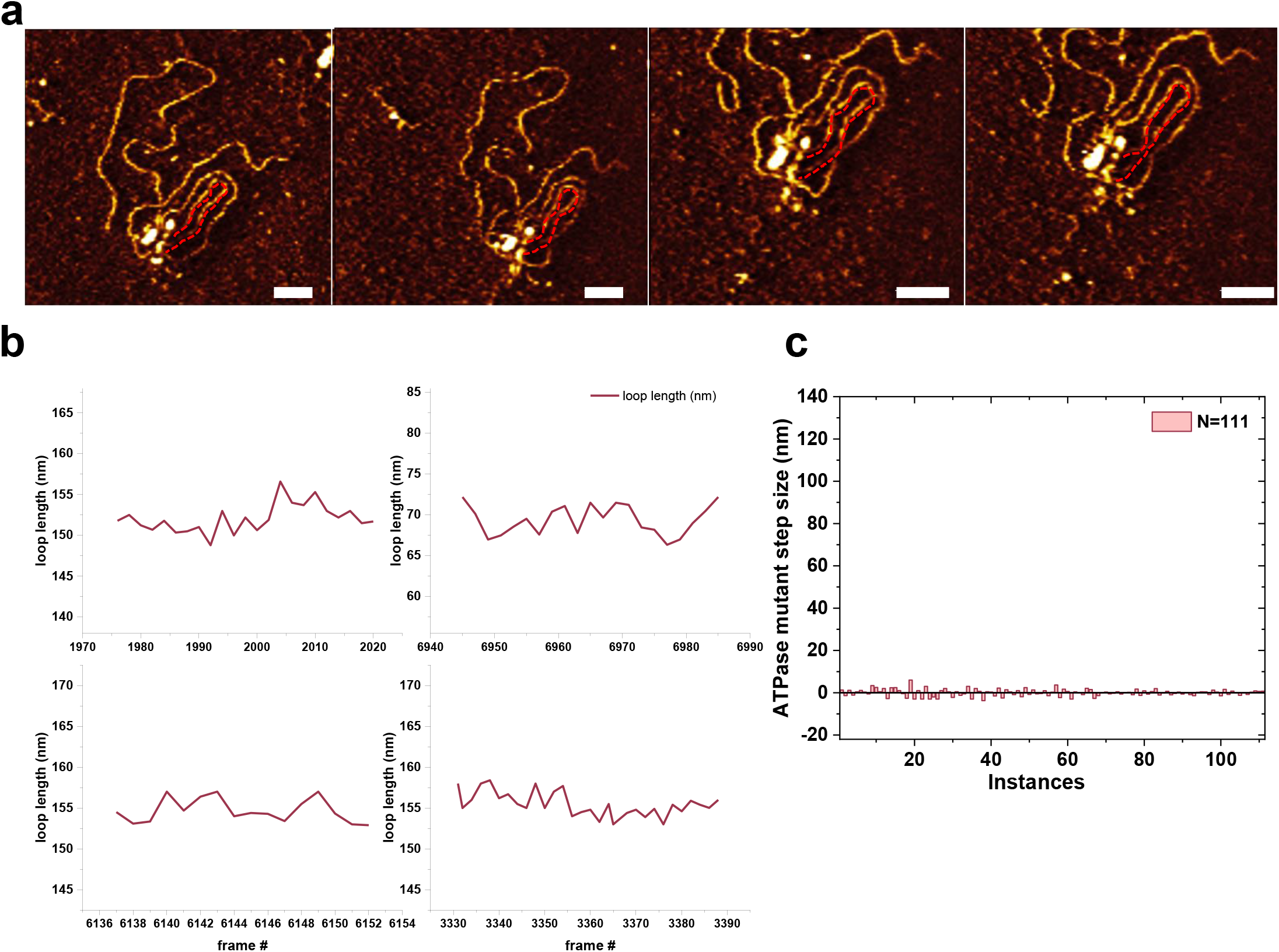
DNA loop length changes mediated by the cohesin^SA1^-NIPBL^c^ ATPase mutant on linear dsDNA. (**a**) Representative time-lapse HS-AFM images of the cohesin^SA1^-NIPBL^c^ ATPase mutant collected at 1 frame/s (+ 4 mM ATP). The average length of the DNA loop was measured at 155.5 nm ± 1.46 (average ± SD) showing no significant change in the length of the DNA loop mediated by the cohesin^SA1^-NIPBL^c^ ATPase mutant. XY scale bar = 50 nm. (**b**) Scatter line plots of the DNA loop length versus the corresponding image frame number for four independent DNA loops mediated by the cohesin^SA1^-NIPBL^c^ ATPase mutant (4 mM ATP). (**c**) Compiled forward (+) and reverse (-) length changes between AFM image frames from 7 independent DNA loops mediated by the cohesin^SA1^-NIPBL^c^ ATPase mutant. Average ± SD for each DNA loop are: 152.1 nm ± 1.7 nm, N=23 frames; 69.4 nm ± 1.8 nm, N=23 frames; 154.7 nm ± 1.5 nm (N=16 frames); 155.5 nm ± 1.5 nm (N=30 frames); 14.4 nm ± 2.4 nm (N=15 frames); 72.3 nm ± 2.9 nm (N=6 frames); and 54.0 nm ± 0.7 nm (N=7 frames).

**Figure S5.**
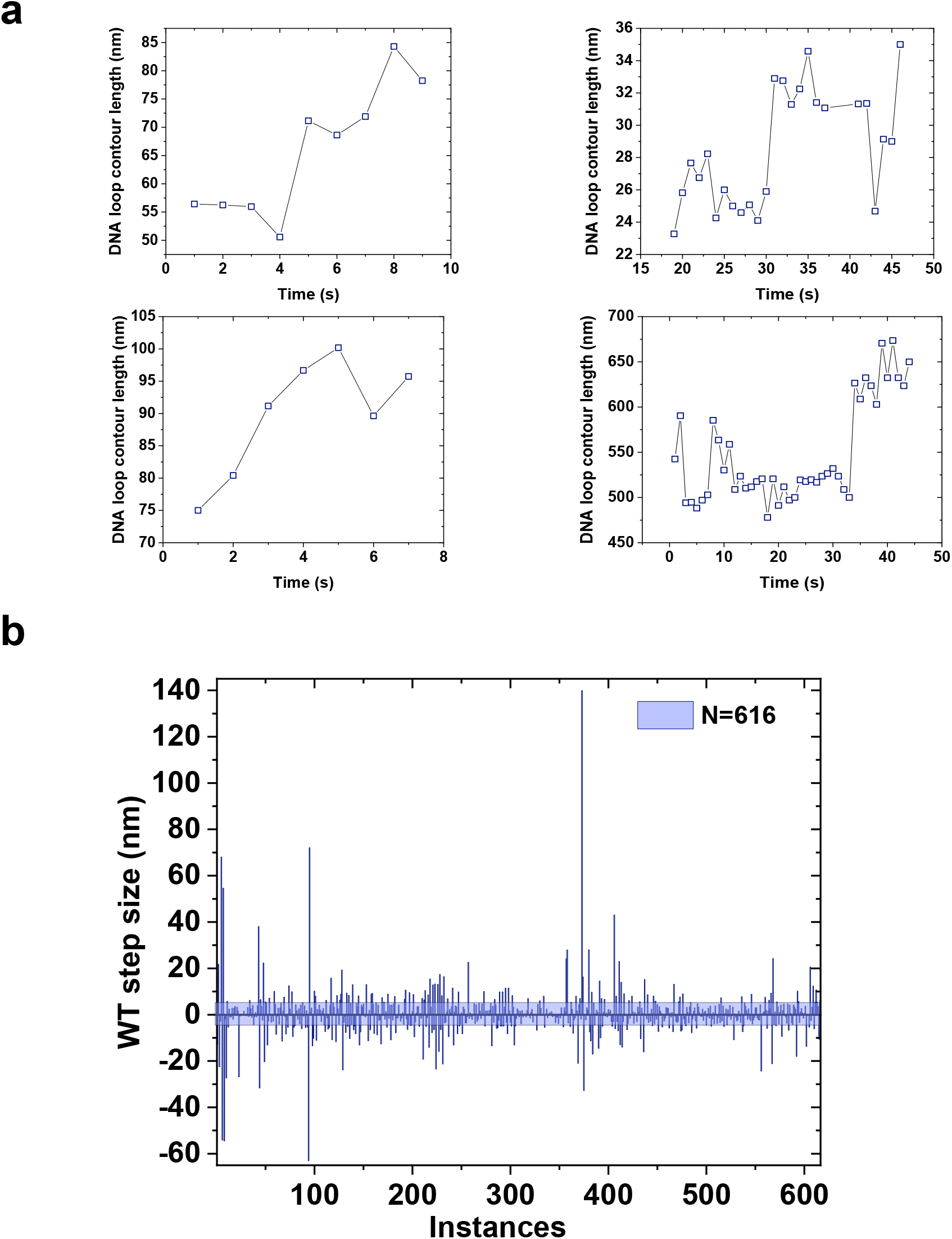
Analysis of DNA loop extrusion by WT cohesin^SA1^-NIPBL^c^ in the presence of ATP. (**a**) Examples of frame-to-frame DNA loop length changes (step sizes per sec) mediated by WT cohesin^SA1^-NIPBL^c^ measured from time-lapse HS-AFM images. (**b**) Compiled DNA looping step sizes mediated by WT cohesin^SA1^-NIPBL^c^ (4 mM ATP) taking into consideration of forward (+: increasing length) and reverse (-: decreasing length) steps. The colored region represents background fluctuations of DNA loop lengths observed when the cohesin^SA1^-NIPBL^c^ ATPase mutant was present (**Figure S4c**).

**Figure S6.**
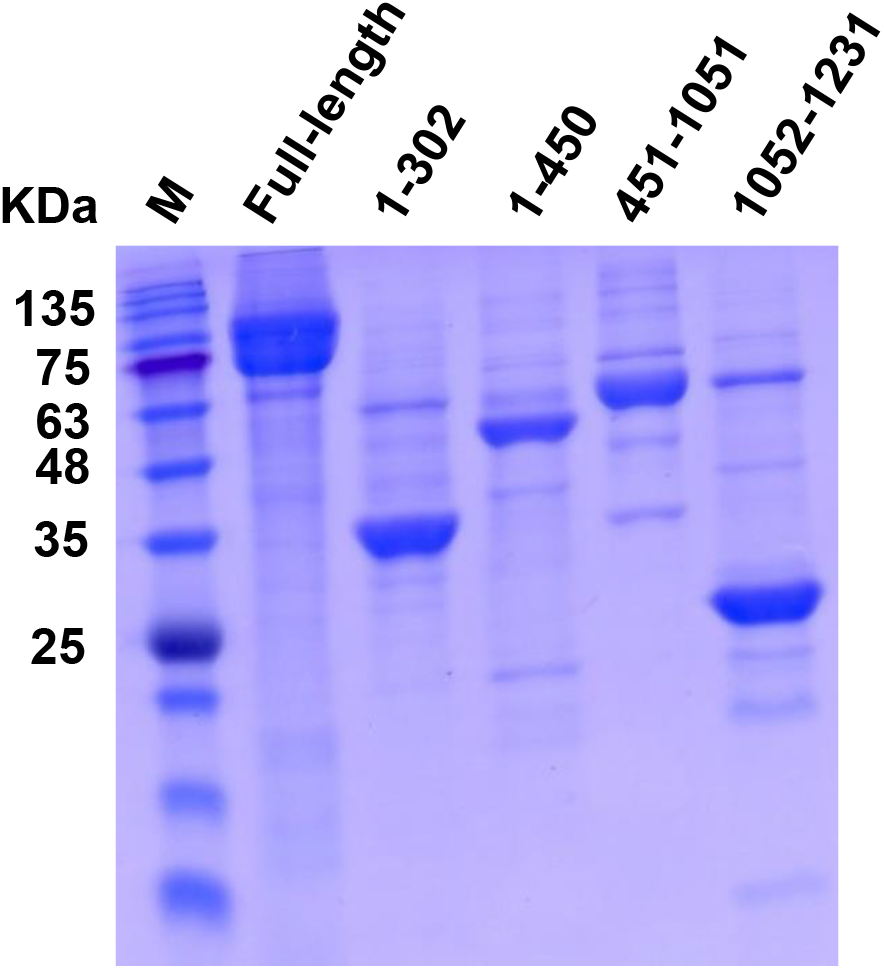
Purification of the full-length SA2 and SA2 fragments. SDS-PAGE of purified WT full-length SA2 1-1231, SA2 1-302, 1-450, 451-1051, and 1052-1231 fragments. M: Molecular weight marker.

**Figure S7.**
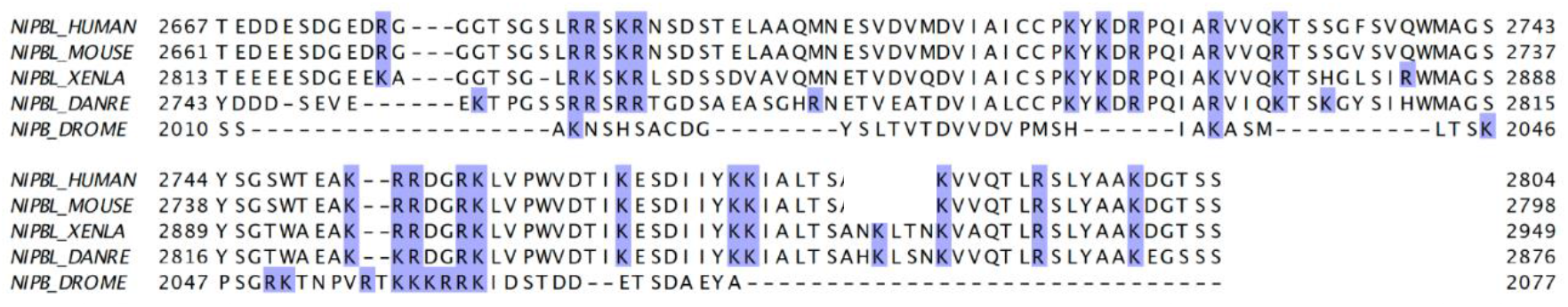
The C-terminal domain of NIPBL contains conserved positively charged residues. Alignment of sequences at the C-terminus of NIPBL from different species.

